# Acute inhibition of heterotrimeric kinesin-2 function reveals mechanisms of intraflagellar transport in mammalian cilia

**DOI:** 10.1101/409318

**Authors:** Martin F. Engelke, Bridget Waas, Sarah E. Kearns, Ayana Suber, Allison Boss, Benjamin L. Allen, Kristen J. Verhey

## Abstract

The trafficking of components within cilia, called intraflagellar transport (IFT), is powered by kinesin-2 and dynein-2 motors. Loss of function in any subunit of the heterotrimeric KIF3A/KIF3B/KAP kinesin-2 motor prevents ciliogenesis in mammalian cells and has hindered an understanding of how kinesin-2 motors function in IFT. We used a chemical-genetic approach to engineer an inhibitable KIF3A/KIF3B (i3A/i3B) kinesin-2 motor that is capable of rescuing WT motor function in *Kif3a/Kif3b* double-knockout cells. Inhibitor addition blocks ciliogenesis or, if added to ciliated cells, blocks IFT within two minutes, which leads to a complete loss of primary cilia within six hours. The kinesin-2 family members KIF3A/KIF3C and KIF17 cannot rescue ciliogenesis in *Kif3a/Kif3b* double-knockout cells nor delay the disassembly of full-formed cilia upon i3A/i3B inhibition. These data suggest that KIF3A/KIF3B/KAP is the sole and essential motor for cilia assembly and function in mammalian cells, indicating a species-specific adaptation of kinesin-2 motors for IFT function.

## INTRODUCTION

Cilia are microtubule-based organelles that protrude from the surface of almost every cell in the body. The beating motion of motile cilia powers cell motility (e.g. sperm) or propels fluids over organ surfaces (e.g. respiratory tract). Primary cilia, in contrast, are largely immotile and function as cellular antennae to sense extracellular stimuli and integrate cellular signaling pathways [1–3]. For example, in vertebrates, Hedgehog (HH) signaling critically depends on the presence of a primary cilium to direct cell proliferation and cell differentiation during development and adult homeostasis [4, 5]. Given the central functions of cilia as motile organelles and cellular signaling hubs, it is not surprising that ciliary malfunction gives rise to a collection of disabling and sometimes life-threatening diseases and syndromes (the ciliopathies) with phenotypes such as developmental malformations, cystic kidney disease, mental retardation, retinal degeneration, and cancer [6, 7].

Intraflagellar transport (IFT) involves the active transport of proteins through the ciliary shaft and is indispensable for the genesis, structural maintenance, and function of all types of cilia [8, 9]. At the base of the cilium, IFT proteins assemble into linear arrays of repeating IFT-A and IFT-B complexes that are named IFT trains based on their beads-on-a-string-like appearance in transmission electron microscopy and tomography images [10–12]. Molecular motor proteins of the kinesin-2 family bind to IFT trains and transport them along the microtubules of the axoneme to the tip of the cilium [13–16], and dynein-2 motors drive transport back to the cell body [17–19].

Current models of IFT are based on genetics and imaging in invertebrates such as *C. elegans* and *Chlamydomonas*, but these models are inadequate to describe kinesin-2 functions in IFT and cilia signaling in mammalian cells. For example, the homodimeric kinesin-2 motor OSM-3 is capable of building a full-length cilium in *C. elegans* but has no homolog in *Chlamydomonas* and its mammalian homolog KIF17 appears to be dispensible for cilia formation in mice [16, 20–24]. On the other hand, the heterotrimeric kinesin-2 motor is dispensable for cilia formation in *C. elegans* but is essential in *Chlamydomonas* and mice [13, 15, 16]. The complete absence of cilia in mammalian cells and organisms upon loss of KIF3A/KIF3B/KAP function has hindered an understanding of the roles of this and other kinesin motors in IFT during ciliary assembly and signaling. Here we use a novel chemical-genetic strategy [25] to engineer an inhibitable KIF3A/KIF3B motor and abruptly inhibit its function in mammalian primary cilia using a highly-specific small molecule.

We find that kinesin-2 inhibition results in a rapid cessation of IFT and complete disassembly of the primary cilium within ∼ 6 hours. This is in contrast to primary cilia that are largely intact after 12 hours of dynein-2 inhibition. Expression of the homodimeric kinesin-2 motor KIF17 neither rescues ciliogenesis nor delays the breakdown of cilia following heterotrimeric kinesin-2 inhibition. These results provide the foundation of a comprehensive model that defines the critical features of IFT during ciliogenesis and ciliary signaling in mammalian cells.

## RESULTS

### Kif3a; Kif3b-deficient NIH-3T3 cells cannot generate primary cilia

To study the function of heterotrimeric kinesin-2 in IFT in mammalian primary cilia, we generated NIH-3T3 cells that lack expression of both heterotrimeric kinesin-2 motor subunits, KIF3A and KIF3B (*Kif3a*^*-/-*^;*Kif3b*^*-/-*^ cells), using CRISPR/Cas9-mediated genome editing [26]. A cell line with a one base pair (bp) insertion and a one bp deletion in the *Kif3a* alleles and a one bp insertion and a 123 bp insertion in the *Kif3b* alleles was generated (**Supp Fig. 1**). All altered open reading frames code for premature stop codons, presumably resulting in nonsense-mediated mRNA decay [27]. Cell cycle arrest induced by serum starvation resulted in the generation of primary cilia in the parental NIH-3T3 cell line transfected with a control (mCherry) plasmid but not in the *Kif3a*^*-/-*^;*Kif3b*^*-/-*^ cells (**Fig. 1a,b,e**). Expression of either KIF3A or KIF3B alone was not sufficient to rescue ciliogenesis, whereas co-expression of KIF3A and KIF3B fully rescued the ability to generate a primary cilium (**Fig. 1c-e**). The ability of ectopic KIF3A and KIF3B expression to rescue the loss of cilia phenotype confirms that the knockout is on target and that there is no expression of a dominant-negative acting kinesin species from the CRISPR/Cas9-modified alleles.

**Figure 1.**
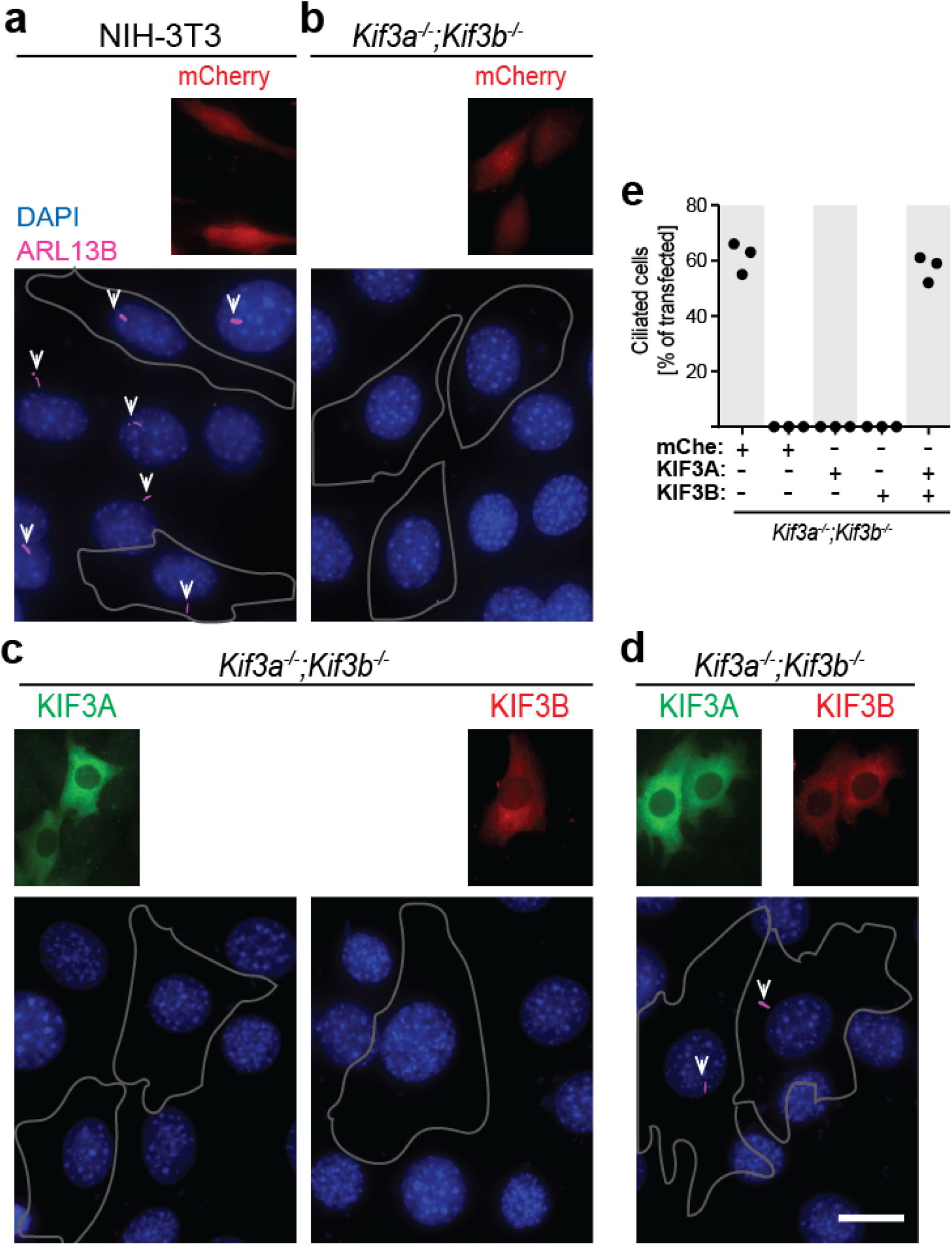
KIF3A/KIF3B-deficient NIH-3T3 cells are not able to generate primary cilia. (**a**) Parental (NIH-3T3) and (**b**-**d**) double knockout (*Kif3a*^*-/-*^;*Kif3b*^*-/-*^) cells ectopically expressing (**a-b**) mCherry or (**c-d**) KIF3A-mNeonGreen and/or KIF3B-mCherry were serum-starved for two days then fixed and stained with DAPI (blue) and an antibody to a marker of the ciliary membrane (ARL13B; magenta). Transfected cells are indicated by a white outline. Primary cilia are marked by a white arrowhead. Scale bar, 10 µm. (**e**) Quantification of the percentage of transfected cells that generate a primary cilium. Each dot represents the average of one independent experiment. n ≥ 94 transfected cells per condition.

### Characterization of inhibitable kinesin-2 motor constructs

We previously described two strategies to generate inhibitable kinesin motors [28]. For generating inhibitable forms of the kinesin-2 motor KIF3A/KIF3B/KAP, we chose to implement the B/B strategy based on the selectivity and applicability of the B/B homodimerizer and the fact that inhibition of KIF3A/KIF3B motors with this strategy will not affect KIF3A/KIF3C motors. The B/B strategy makes use of the ability of the small molecule B/B homodimerizer to rapidly and specifically mediate the dimerization of DmrB domains [29, 30]. By fusing the DmrB domains to the N-termini of KIF3A and KIF3B, the B/B homodimerizer will crosslink and there by inhibit the stepping motion of the heterotrimeric kinesin-2 motor in a B/B (inhibitor)-dependent fashion (**Fig. 2a**). Inhibitable KIF3A and KIF3B constructs are referred to as i3A and i3B, respectively.

**Figure 2.**
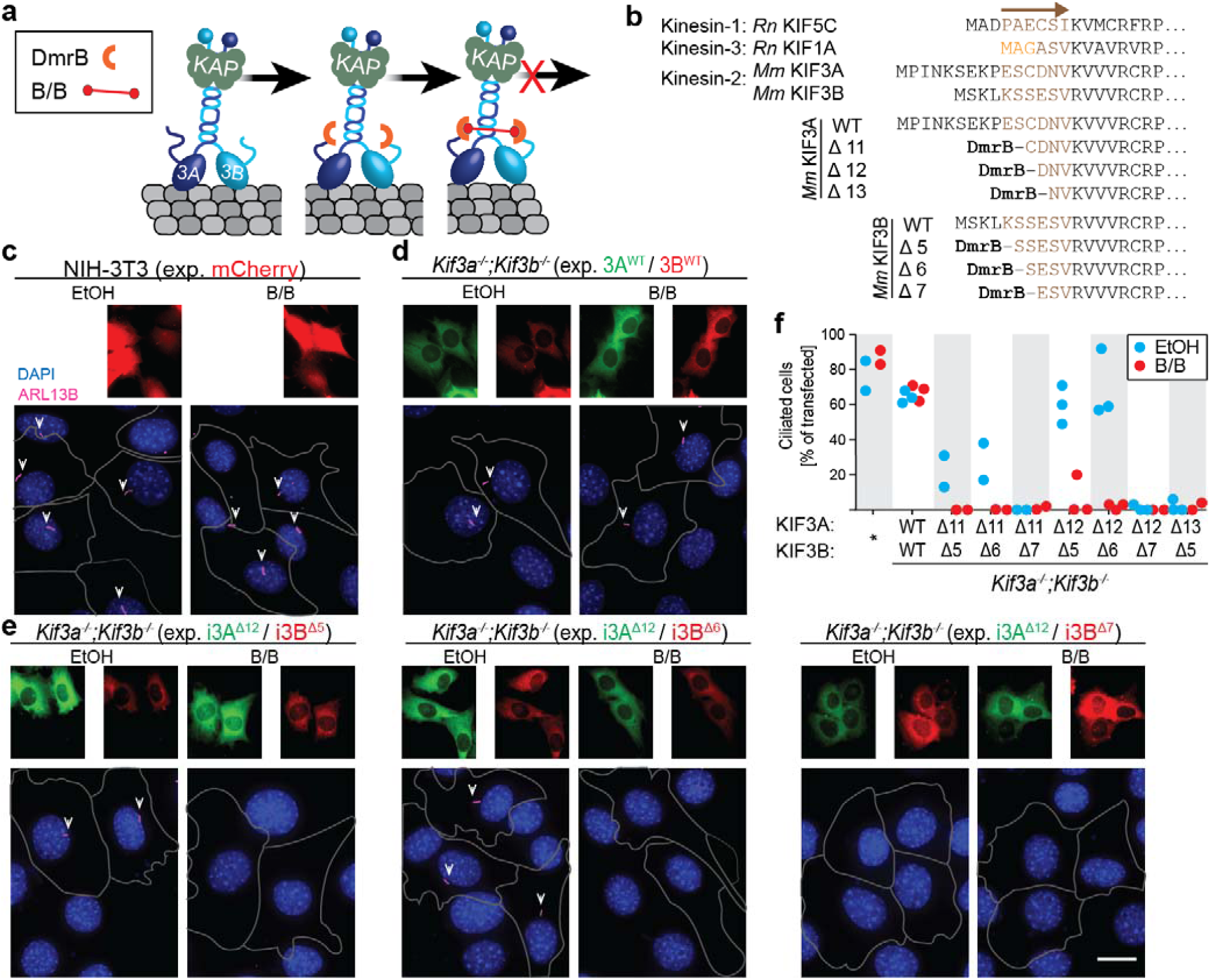
Characterization of inhibitable KIF3A/KIF3B motors in ciliogenesis. (**a**) Engineering strategy to generate inhibitable KIF3A/KIF3B motors. (left) The WT heterotrimeric kinesin-2 motor (KIF3A, dark blue; KIF3B, light blue; KAP3, green) moves along microtubules (gray) and transports molecular cargo (not shown). (middle) The inhibitable motor comprises KIF3A and KIF3B subunits engineered with a Dmr-B domain (orange semicircle) fused to their N-termini. This motor moves along microtubules and transports cargo similar to the WT motor. (right) Addition of the B/B homodimerizer (red dumbbell) crosslinks the motor domains and thereby inhibits processive motor motility. (**b**) Alignment of the N-terminal residues of kinesin-1 (KIF5C), kinesin-3 (KIF1A), and kinesin-2 (KIF3A, KIF3B) (top). The cover strand (brown arrow and text) is the most N-terminal structural element of the motor domain. Orange text indicates residues that were truncated in KIF1A to generate an inhibitable motor [25]. To generate inhibitable kinesin-2 motors, the position of the DmrB-domain relative to the cover strand was modified by truncation of 11, 12, or 13 resides of KIF3A (middle) or truncation of 5, 6, or 7 residues of KIF3B (bottom). (**c-e**) Representative images of (**c**) parental NIH-3T3 cells expressing mCherry control or (**d-e**) *Kif3a*^*-/-*^;*Kif3b*^*-/-*^ cells expressing (**d**) WT or (**e**) inhibitable kinesin-2 motors [KIF3A constructs tagged with mNG (green), KIF3B constructs tagged with mCherry (red)]. The cells were treated with ethanol (vehicle control) or B/B, serum-starved for two days, then fixed and stained with DAPI (blue) and an antibody against ARL13B (magenta). Transfected cells are indicated by a white outline. Primary cilia are marked by a white arrowhead. Scale bar, 10 µm. (**f**) Quantification of the percentage of transfected cells that generate a primary cilium. (*) NIN-3T3 cells transfected with mCherry control. Each spot represents the mean of one independent experiment. n ≥ 57 transfected cells per condition.

As a starting point for the optimization of engineering inhibitable motors, we fused the DmrB domains directly to the N-termini of the KIF3A and KIF3B motor domains, since this DmrB fusion site had yielded inhibitable motors for kinesin-1 [25]. Transfection of the resulting DmrB-KIF3A and DmrB-KIF3B fusion constructs rescued ciliogenesis in *Kif3a*^*-/-*^;*Kif3b*^*-/-*^ cells, but the addition of the B/B inhibitor had no effect on ciliogenesis (data not shown), suggesting that these are active motors but are not inhibitable. We thus considered that engineering of the kinesin-3 motor KIF1A revealed that only one DmrB-insertion site yielded a functional, inhibitable motor [25]. To find an analogous insertion site for the kinesin-2 motor subunits, we compared the N-terminal amino acid sequences of kinesin-1, -2, and -3 motors and designed Dmr-B insertion sites for KIF3A and KIF3B comparable to that of the inhibitable kinesin-3 KIF1A motor (**Fig. 2b**). We created a series of fusion constructs in which the DmrB domain was fused to the N-terminus of KIF3A truncated by 11 (i3A^Δ11^), 12 (i3A^Δ12^), or 13 (i3A^Δ13^) amino acids. In similar fashion, we fused the DmrB domain to the N-terminus of KIF3B truncated by five (i3B^Δ5^), six (i3B^Δ6^), or seven (i3B^Δ7^) amino acids (**Fig. 2b**). We compared the ability of each i3A construct to pair with each i3B construct and generate primary cilia in the absence but not in the presence of B/B inhibitor. Fusion of the DmrB domain to the Δ13 truncation of KIF3A (i3A^Δ13^) or to the Δ7 truncation of KIF3B (i3B^Δ7^) resulted in a failure to generate primary cilia (i3A^Δ13^/i3B^Δ5^, i3A^Δ11^/i3B^Δ7^, and i3A^Δ12^/i3B^Δ7^ in **Fig. 2e-f** and data not shown), indicating that engineering the i3A^Δ13^ and i3B^Δ7^ constructs resulted in non-functional motors. Fusion of the DmrB domain to the Δ12 truncation of KIF3A (i3A^Δ12^) resulted in functional motors when paired with the i3B^Δ5^ or i3B^Δ6^ constructs of KIF3B as expression of i3A^Δ12^/i3B^Δ5^ or i3A^Δ12^/i3B^Δ6^ rescued ciliogenesis in *Kif3a*^*-/-*^;*Kif3b*^*-/-*^ cells to a similar extent as expression of WT KIF3A/KF3B (i3A^Δ11^/i3B^Δ5^, i3A^Δ11^/i3B^Δ6^ and WT/WT in **Fig. 2e-f**). Importantly, addition of B/B inhibitor completely blocked ciliogenesis in i3A^Δ12^/i3B^Δ6^-expressing cells and to a lesser extent in i3A^Δ12^/i3B^Δ5^-expressing cells (**Fig. 2e-f**). Fusion of the DmrB domain to the Δ11 truncation of KIF3A (i3A^Δ11^) also rescued ciliogenesis when paired with the i3B^Δ5^ or i3B^Δ6^ constructs of KIF3B, albeit at a lower frequency than cells expressing the WT motors, and their function was blocked by B/B inhibitor (**Fig. 2f**). Because the i3A^Δ12^/i3B^Δ6^ combination was fully functional for ciliogenesis and exhibited the best inhibition, this combination was utilized throughout the remainder of this work to selectively block heterotrimeric kinesin-2 function. The inhibition of i3A^Δ12^/i3B^Δ6^ was sensitive to B/B inhibitor concentration in dose-responsive manner (**Supp Fig. 2**).

We next tested whether primary cilia generated upon expression of inhibitable i3A^Δ12^/i3B^Δ6^ motors are competent to mediate Hedgehog (HH) signaling. We first tested HH pathway activation in response to Smoothened agonist (SAG) using a reporter construct in which luciferase expression is driven by a HH-responsive promoter [31]. Addition of SAG resulted in a ∼4-fold induction in luciferase activity for *Kif3a*^*-/-*^;*Kif3b*^*-/-*^ cells expressing WT KIF3A/KIF3B or the inhibitable i3A^Δ12^/i3B^Δ6^ motor (**Fig. 3b**), indicating that the inhibitable motor generates fully HH-responsive cilia. Addition of the B/B inhibitor diminished the HH response to SAG treatment in *Kif3a*^*-/-*^;*Kif3b*^*-/-*^ cells expressing the i3A^Δ12^/i3B^Δ6^ motor but not in cells expressing the WT KIF3A/KIF3B motor (**Fig. 3b**). As a second approach to test the HH response in cells expressing the inhibitable kinesin-2, we used a constitutively-active version of the HH transcription factor GLI2 (GLI2ΔN) [32]. Previous work suggested that expression of GLI2ΔN ectopically activates HH target gene expression, and that a loss of primary cilia can result in greater pathway activation in this context [33]. Expression of GLI2ΔN in *Kif3a*^*-/-*^;*Kif3b*^*-/-*^ cells expressing WT KIF3A/KIF3B or the inhibitable i3A^Δ12^/i3B^Δ6^ motor resulted in ∼200-fold luciferase induction (**Fig. 3c**). Addition of B/B inhibitor resulted in a greater increase in luciferase activity induction only in cells expressing the i3A^Δ12^/i3B^Δ6^ motor, reflecting pathway hyper-activation only in cells that lack a functional primary cilium (**Fig. 3c**).

**Figure 3.**
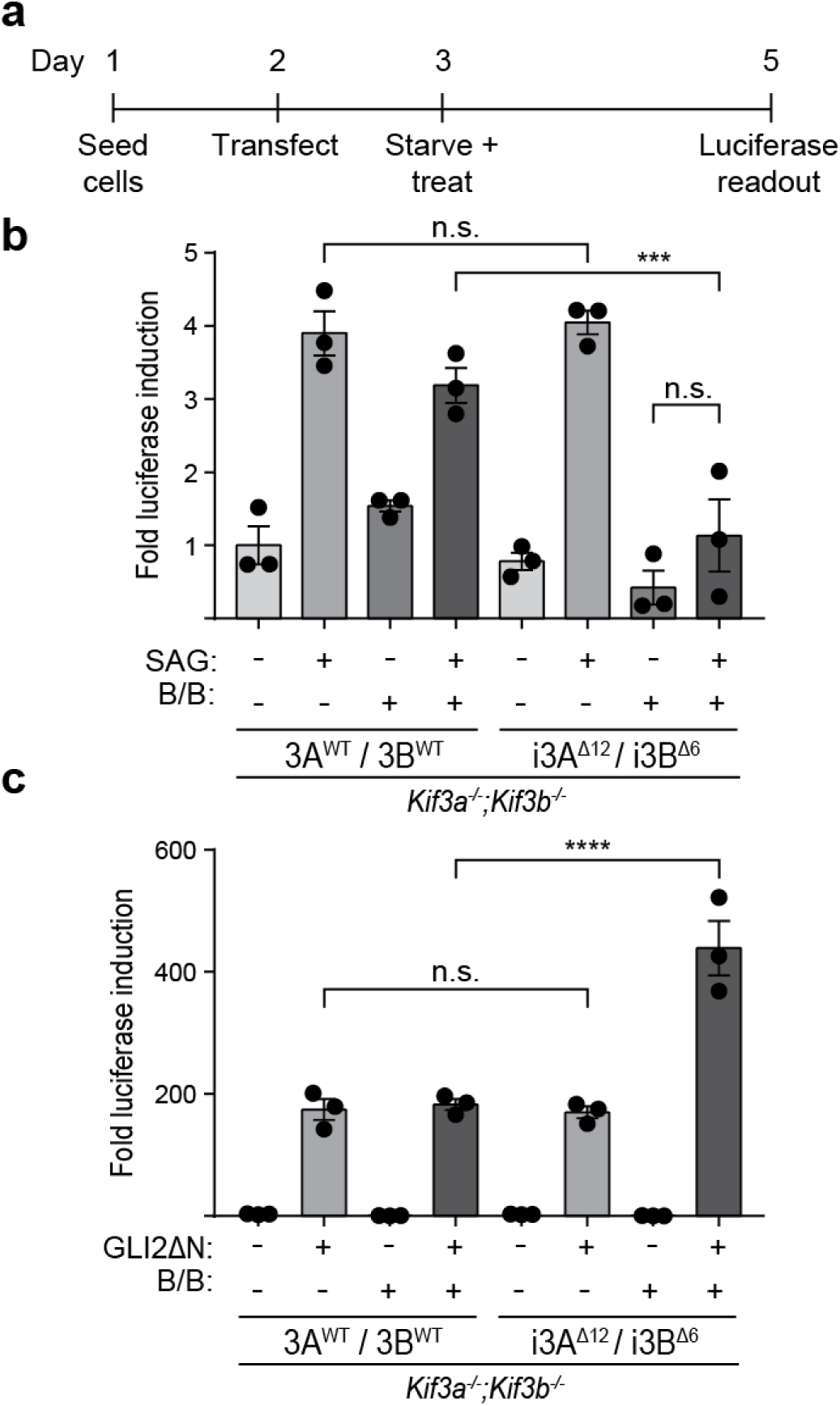
Inhibitable kinesin-2 motors generate hedgehog (HH)-responsive cilia. (**a-c**) HH signaling assays. **(a)** Schematic of the experimental setup. Cells were seeded on day 1 and transfected on day 2 with plasmids for expressing a HH-responsive luciferase plasmid and a control beta-galactosidase plasmid together with **(b)** WT or inhibitable KIF3A/KIF3B or (**c**) GLI2ΔN and WT or inhibitable KIF3A/KIF3B. The following day, the cells were serum-starved and treated with Sonic HH agonist (SAG) and/or B/B inhibitor. Two days later, luciferase activity was measured and normalized to beta-galactosidase activity. (**b**) Quantification of HH signaling in response to SAG. No significant difference (n.s.) was observed in luciferase induction between cells expressing WT or inhibitable motors. Cells expressing inhibitable KIF3A/KIF3B and treated with B/B inhibitor showed a significant decrease in response to SAG (***p = 0.0001, Sidak’s multiple comparisons test) as compared to cells expressing WT KIF3A/KIF3B and treated with SAG. (**c**) Quantification of HH signaling in response to GLI2ΔN expression. No significant difference (n.s.) was observed in luciferase induction between cells expressing WT or inhibitable motors. Cells expressing inhibitable KIF3A/KIF3B and treated with B/B inhibitor showed a significant increase in GLI2ΔN-induced luciferase expression (****p < 0.0001, Sidak’s multiple comparisons test) as compared to cells expressing WT KIF3A/KIF3B. Error bars indicate SEM.

In summary, we find that expression of the i3A^Δ12^ and i3B^Δ6^ constructs in *Kif3a*^*-/-*^;*Kif3b*^*-/-*^ cells results in a *bona fide* inhibitable kinesin-2 motor. The engineered i3A^Δ12^/i3B^Δ6^ motor is referred to as i3A/i3B throughout the rest of the manuscript.

### Inhibition of KIF3A/KIF3B results in the stalling of IFT trains and their exclusion from cilia

The generation of an inhibitable kinesin-2 motor enables us, for the first time, to directly examine the role of KIF3A/KIF3B/KAP motors during IFT in fully-formed cilia. To investigate this, *Kif3a*^*-/-*^;*Kif3b*^*-/-*^ cells were transfected with plasmids for expressing the inhibitable i3A/i3 motor together with a fluorescently-tagged subunit of the IFT-B complex (IFT88-mNG; mNeonGreen) as in previous studies [34, 35]. Analysis of kymographs generated from live-cell imaging experiments revealed that IFT88-marked IFT trains moved processively towards the tip of the cilium, paused for variable durations, and then trafficked back towards the base (**Fig. 4a**), similar to what has been observed previously [34, 35]. Anterograde and retrograde speeds were on the order of 0.7 μm/s, consistent with previously measured IFT velocities in mammalian cilia [35–37]. To determine the impact of inhibition of KIF3A/KIF3B function on IFT, cilia expressing the inhibitable motor and IFT88-mNG were imaged and then B/B inhibitor, or ethanol vehicle as a control, was added and imaging of the same cilium was resumed. IFT motion persisted after treatment with the vehicle (ethanol), however, no IFT motion was observed in the anterograde or retrograde direction within five min of treatment with the B/B inhibitor (**Fig. 4a**). In the example shown in **Supp Fig. 3**, three IFT trains were observed to move within the cilium before all transport ceased and no more IFT trains entered the cilium within two minutes of B/B inhibitor addition.

**Figure 4.**
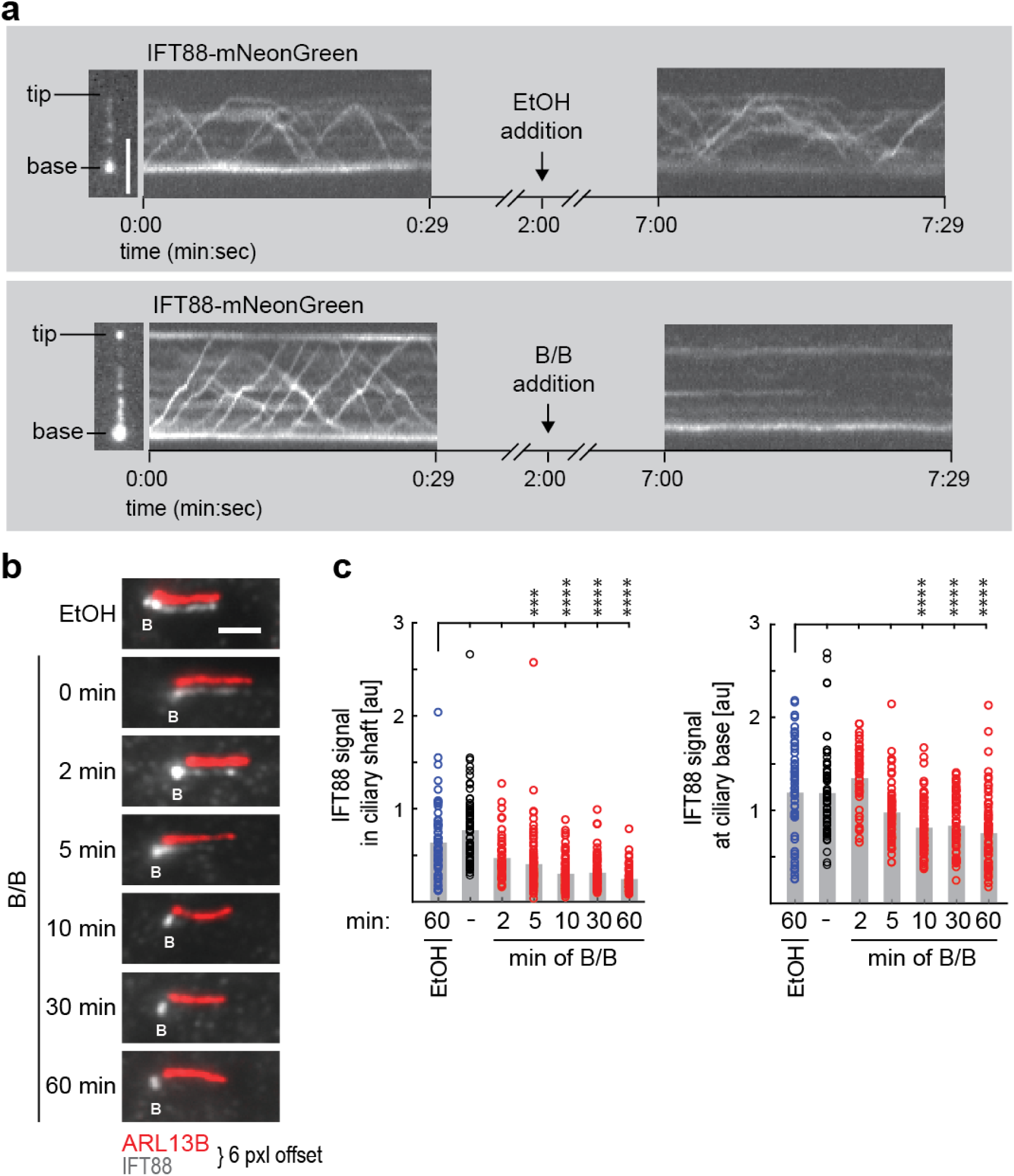
Inhibition of heterotrimeric kinesin-2 stops IFT and results in a complete loss of IFT trains from cilia. (**a**) Live-cell imaging of *Kif3a*^*-/-*^;*Kif3b*^*-/-*^ cells expressing inhibitable kinesin-2 (i3A^Δ12^ and i3B^Δ6^) motors and IFT88-mNG. An image of a representative cilium (left) and corresponding kymographs before treatment (middle) and after addition of ethanol vehicle (EtOH, top right) or B/B inhibitor (B/B, bottom right) are shown. Imaging frequency, 5 Hz; scale bar, 2 µm. (**b-c**) *Kif3a*^*-/-*^;*Kif3b*^*-/-*^ cells transfected with inhibitable kinesin-2 (i3A^Δ12^ and i3B^Δ6^) motors were serum-starved, treated with ethanol vehicle or B/B inhibitor for the indicated times, and then fixed and stained with antibodies to the IFT protein IFT88, the basal body marker glutamylated tubulin, and the ciliary marker ARL13B. (**b**) Representative images of IFT88 (gray) and ARL13B (red) in the primary cilium with the fluorescence signals offset by 6 pixels for clarity. B indicates the ciliary base. Scale bar, 2 µm. (**c**) Quantification of the average fluorescence intensity of IFT88 in the ciliary shaft (left) and at the ciliary base (right). Colored circles indicate the average fluorescence intensity per cell and gray bars show the average fluorescence intensity of all cells in that condition. *** p = 0.0001 and **** p < 0.0001, Dunn’s test.

To examine the effects of kinesin-2 inhibition across a large number of cells, we fixed and stained cells at various time points after B/B treatment. A significant loss of IFT88 from the shaft of the primary cilium was observed within five minutes of B/B treatment and no further loss of IFT88 signal in the cilium was detected after 10 minutes of treatment (**Fig. 4b-c**). Thus, blocking kinesin-2 function leads to a complete absence of IFT88-marked IFT trains moving within the primary cilium. In contrast, we observed an increase in IFT88 signal at the base of the cilium within the first two minutes after B/B treatment. However, this increase was transient and not statistically significant as the IFT88 signal dropped below baseline conditions with longer B/B treatment (**Fig. 4b-c**). Taken together, these data show that all IFT motions in cells expressing the i3A/i3B motor are blocked within two minutes of B/B inhibitor addition, leading to a subsequent loss of IFT from the primary cilium.

### KIF3A/KIF3B-mediated IFT is essential for the maintenance of ciliary structure

How does an acute and complete block in IFT affect the stability of the primary cilium? To address this question, cells expressing the i3A/i3B motor were allowed to generate a primary cilium and then the effects of B/B addition on the length and abundance of cilia were quantitatively assessed (**Fig. 5a**). Treatment with B/B resulted in a decrease in the percentage of cells with a primary cilium as early as one hour after B/B treatment (63% ciliated vs 83% in vehicle-treated). The percentage of ciliated cells continued to decline throughout an eight-hour time course until only 2% of cells expressing the i3A/i3B motor retained a primary cilium after B/B treatment (**Fig. 5b**). For the cilia that remained after B/B treatment, there was also a decrease in the average ciliary length as measured by both the ciliary membrane marker ARL13B and the axonemal marker acetylated α-tubulin. The apparent loss in ciliary length was slow and continuous with the average cilium length decreasing from 2.6 μm in vehicle-treated cells (ethanol) to 1.7 μm six hours after B/B treatment (**Fig. 5c**). Although the average ciliary length increased at eight hours compared to control-treated cells, this value is likely an outlier as only one cilium persisted at this time point.

**Figure 5.**
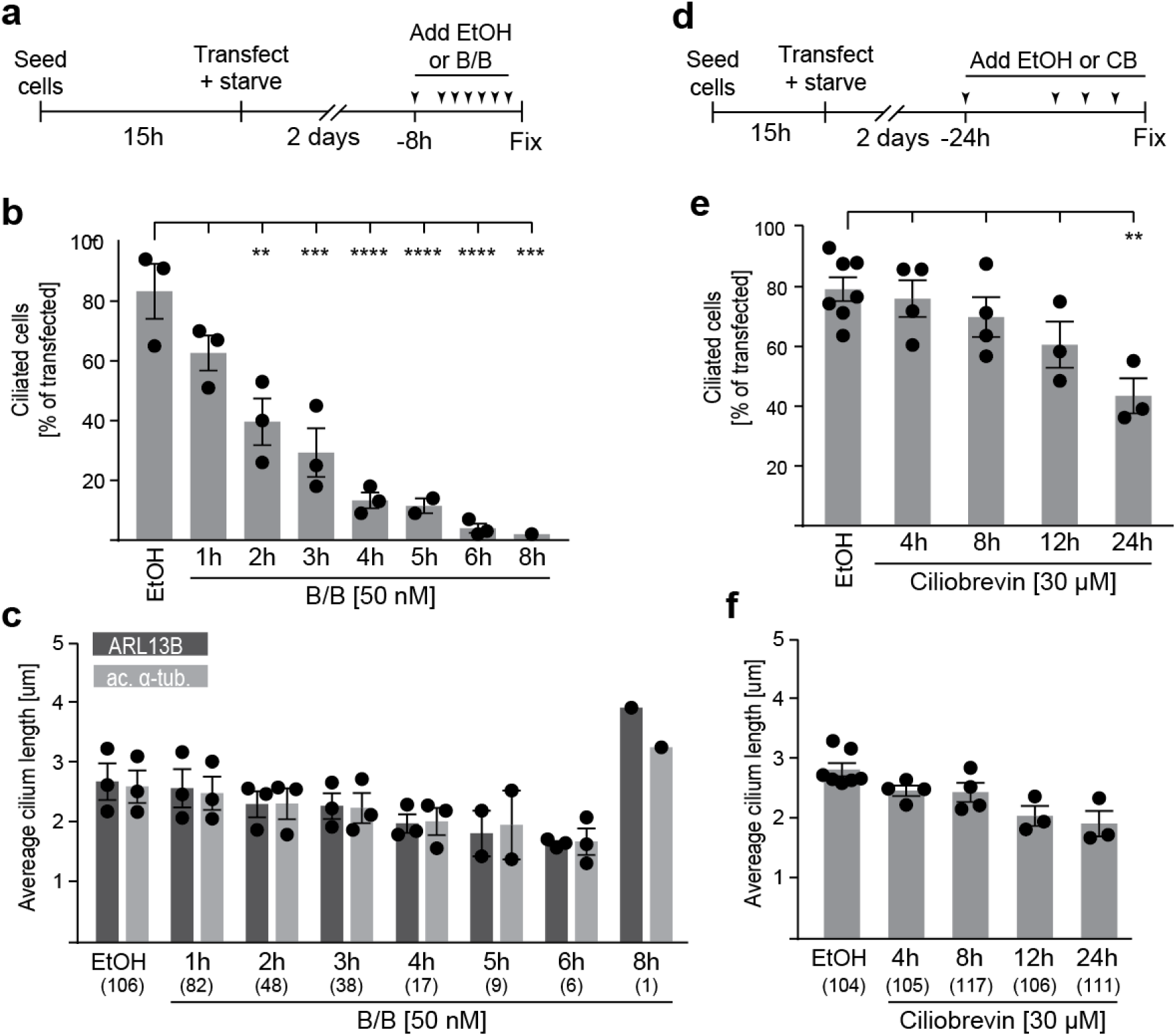
Inhibition of heterotrimeric kinesin-2 results in the shortening and disassembly of cilia. **(a-c)** Inhibition of heterotrimeric kinesin-2. (**a**) Schematic of experimental setup. *Kif3a*^*-/-*^;*Kif3b*^*-/-*^ cells were serum-starved and transfected with inhibitable kinesin-2 (i3A^Δ12^ and i3B^Δ6^) motors. The cells were treated with ethanol (control) or B/B inhibitor at the indicated time points and then fixed and stained with antibodies against the axonemal marker acetylated α-tubulin and the ciliary membrane marker ARL13B. (**b,c**) Quantification of the (**b**) percentage of cells that generate a primary cilium and (**c**) cilium length after treatment with B/B inhibitor for the indicated times. Each spot indicates the mean of one independent experiment. N values are indicated in parenthesis on the x-axis of (**c**). Error bars, SEM. ** p = 0.002; *** p < 0.0004; **** p < 0.0001 as determined by post hoc Sidak’s test. (**d-f**) Inhibition of cytoplasmic dynein. **(d)** Schematic of experimental setup. The cells were treated with ethanol (control) or ciliobrevin (CB) at the indicated time points. (**e-f**) Quantification of the (**e**) percentage of cells with a primary cilium and (**f**) cilium length after treatment with Ciliobrevin D for the indicated times. N values are indicated in parenthesis on the x-axis of (**f**). ** p = 0.0016, Dunnett’s test.

To compare cilium disassembly in response to inhibition of anterograde versus retrograde transport, we treated cells with Ciliobrevin D, an inhibitor of the retrograde IFT motor dynein-2 [38] (**Fig. 5d**). Little to no change in the percentage of ciliated cells was determined after eight hours of Ciliobrevin D treatment and only after 24 hours of Ciliobrevin D treatment was a significant decrease in ciliated cells observed (43% ciliated versus 79% in the ethanol control) (**Fig. 5e**). In the same time frame, the average cilia length dropped from 2.8 μm in ethanol-treated cells to 1.9 μm in Ciliobrevin D-treated cells (**Fig. 5f**). These results indicate that continual anterograde IFT is critical for the maintenance of the structure of cilia whereas retrograde IFT is less important for ciliary maintenance.

### KIF17 is not involved in ciliogenesis or cilia maintenance

In addition to KIF3A/KIF3B/KAP, two other motor proteins of the kinesin-2 family, heterotrimeric KIF3A/KIF3C/KAP and homodimeric KIF17, are implicated in driving transport events within mammalian cilia [39–41]. We thus probed an involvement of these motors in ciliogenesis by ectopically expressing either KIF3A/KIF3C or KIF17 in *Kif3a*^*-/-*^;*Kif3b*^*-/-*^ cells. We found that *Kif3a*^*-/-*^;*Kif3b*^*-/-*^ cells expressing either motor for two days were not able to generate cilia (**Fig. 6a**), demonstrating that KIF3A/KIF3B/KAP is the essential motor for ciliogenesis in NIH-3T3 cells. This is consistent with studies demonstrating that loss of function in KIF3A, KIF3B, or KAP results in a complete absence of cilia in vertebrate organisms [15, 42–44]. However, KIF17 binds to IFT-B particles and co-migrates with IFT trains in mammalian cilia [45, 46], suggesting that this motor could take over from KIF3A/KIF3B/KAP and function in cilia maintenance in a similar manner as its worm homolog OSM-3. We tested this hypothesis by co-expressing i3A/i3B and KIF17 in the *Kif3a*^*-/-*^;*Kif3b*^*-/-*^ cells. After cells were fully ciliated, the B/B inhibitor was added to block KIF3A/KIF3B/KAP function and the percentage of ciliated cells and the cilium length was measured. Across a six hour time frame, cells expressing the KIF17 motor displayed an identical loss of cilia and decrease in ciliary length as the cells expressing only the i3A/i3B motor (**Fig. 6c-d**). The observed difference in cilium length at the six-hour time point is likely an outlier, as only one cilium persisted in KIF17-expressing cells after six hours of B/B treatment. These results demonstrate that despite the ability of KIF17 to bind to and co-traffic with IFT trains, KIF17 does not power IFT in cilia of mammalian cells.

**Figure 6.**
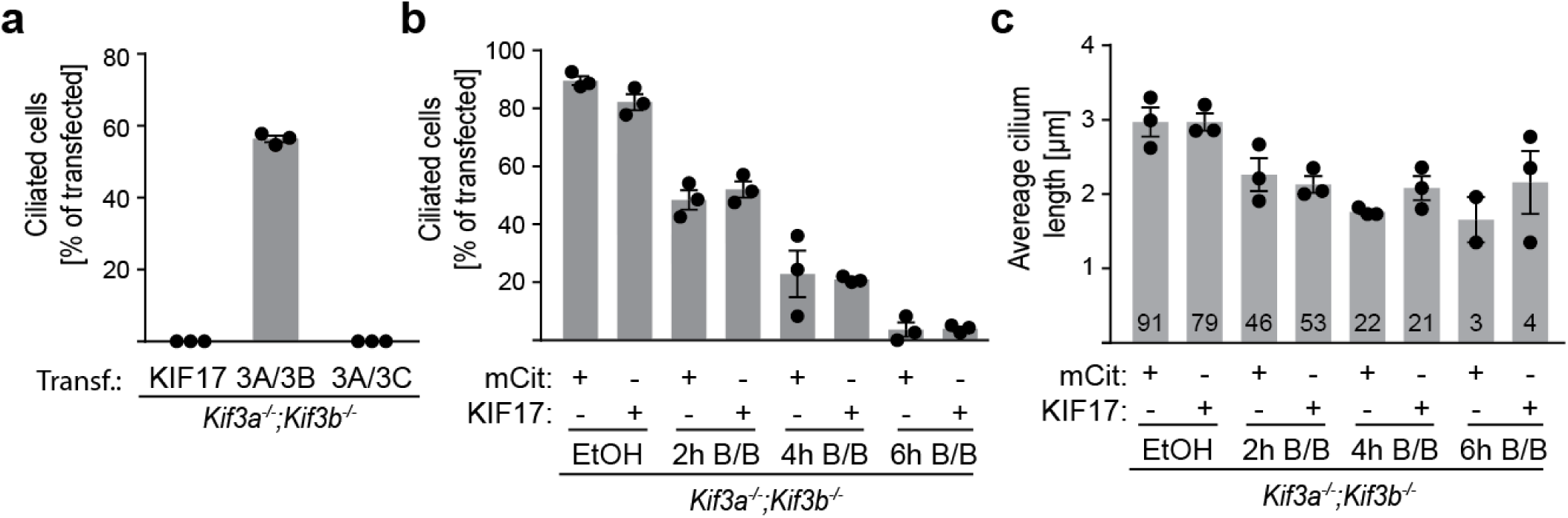
Neither the heterotrimeric KIF3A/KIF3C/KAP motor nor the homodimeric KIF17 can substitute for KIF3A/KIF3B/KAP in mammalian cilia. (**a**) Test for rescue of ciliogenesis. *Kif3a*^*-/-*^;*Kif3b*^*-/-*^ cells were transfected with KIF17, KIF3A and KIF3B (3A/3B), or KIF3A and KIF3C (3A/3C) and serum-starved to allow for ciliogenesis. The cells were fixed and stained with an antibody to the ciliary membrane marker ARL13B and the percentage of transfected cells with a primary cilium was quantified. (**b-c**) Test for rescue of cilia maintenance. *Kif3a*^*-/-*^;*Kif3b*^*-/-*^ cells were co-transfected with inhibitable i3A^Δ12^/i3B^Δ6^ and KIF17 motors and serum-starved. After two days, the cells were treated with ethanol control (EtOH) or B/B inhibitor for 2, 4, or 6 hours and then fixed and stained. The (**b**) percentage of cells with a primary cilium and the (**c**) length of the primary cilium were quantified using the ciliary membrane marker ARL13B. Error bars, SEM. N values are indicated on each bar in (**c**). At each time point, there was no significant difference in the percentage of ciliated cells or in the average cilium length when comparing cells expressing KIF17 and the mCit control. For both conditions (mCit and KIF17), there was a significant decrease in the percentage of ciliated cells at each time point compared to the control ethanol treatment (p<0.0001, Sidak’s test).

## DISCUSSION

### Delineating cellular processes with inhibitable KIF3A/KIF3B/KAP motors

We engineered an inhibitable version of the heterotrimeric KIF3A/KIF3B/KAP kinesin-2 motor that allows, for the first time, specific inhibition of this motor and delineation of its role in IFT in primary cilia. The successful engineering of inhibitable motors from the kinesin-1, kinesin-3 [25], and now kinesin-2 families corroborates the notion that our chemical-genetic approach can be used to generate inhibitable motors across the kinesin family, and likely other motor proteins of interest, without the need for labor-intensive small molecule screens. The inhibition of the engineered motors is highly specific, as it relies on the well-characterized B/B homodimerizer [29, 30], which has no endogenous cellular targets, thus avoiding off-target effects. The inhibition is also rapid, occurring on the timescale of the motor transport functions; this is in contrast to inhibition of transport function for a temperature-sensitive allele of kinesin-2 (fla10-1^ts^) in *Chlamydomonas* where a full block to IFT is not observed until ∼60 min after shifting cells to the non-permissive temperature [13, 14, 47].

The engineered i3A/i3B/KAP motor is fully functional as it restores ciliogenesis in Kif3a^-/-^;Kif3b^-/-^ cells and generates HH-responsive cilia, an important criterion that has not been demonstrated for engineered motors generated by other chemical-genetic approaches [48]. An inability to fully replicate wild-type protein function has also been noted for the fla10-1^ts^ *Chlamydomonas* temperature-sensitive allele as fewer IFT trains are observed in cilia grown at the permissive temperature than in the WT strain [49]. While we have tested our i3A/i3B motor extensively in *Kif3a*^*-/-*^;*Kif3b*^*-/-*^ cells, future work will show whether this motor indeed fully complements the WT protein function in an i3A/i3B mouse model. Importantly, fusion of DmrB domains to both motor-containing subunits, KIF3A and KIF3B, has an added advantage in that it allows a separation of the functions of KIF3A/KIF3B/KAP from those of KIF3A/KIF3C/KAP.

### Inhibition of KIF3A/KIF3B/KAP results in a complete loss of IFT

We observed a rapid block to IFT upon inhibition of the i3A/i3B/KAP motor in mammalian primary cilia. In the absence of B/B inhibitor, we find that IFT88 moves processively along the length of the cilium in anterograde and retrograde directions, consistent with previous work in mammalian cells [45, 50]. Addition of the B/B inhibitor resulted in a rapid block to IFT (within two minutes). In the example in **Supp Fig 3**, three IFT trains can be observed moving in the anterograde direction after the addition of the inhibitor. These trains are likely driven by i3A/i3B/KAP motors that have not yet been inhibited and thus undergo the normal process of being remodeled at the cilium tip and removed from the cilium by retrograde IFT driven by dynein-2.

The most dramatic effect of i3A/i3B inhibition is a rapid block to new IFT trains entering the cilium. One possibility is that B/B inhibitor crosslinks and stops i3A/i3B/KPA motors once they are activated by cargo binding at the base of the cilium. It is also possible that B/B inhibitor crosslinks the motor domains of soluble and autoinhibited kinesin-2 motors, thereby depleting the pool of active kinesin-2 that can be recruited to IFT trains at the base of the cilium. These possibilities are not mutually exclusive and indeed are both compatible with recent work in *Chlamydomonas* demonstrating that IFT trains are assembled at the basal body and kinesin-2 is recruited to the train just before it is injected into the cilium [12]. Further work will examine the ability of B/B inhibitor to block kinesin-2 transport and/or deplete the soluble pool of autoinhibited motors.

Also consistent with these possibilities is the transient increase in IFT88 intensity at the base of the cilium after B/B inhibitor addition. Although this transient increase was not statistically significant, it is compatible with the idea that IFT particles assemble at the base of the cilium and are then loaded on a kinesin-2 motor for import into the cilium and IFT. That the increase in IFT88 is transient and moderate indicates that the number of IFT binding sites at the base of the cilium is limited, consistent with the recent work in *Chlamydomonas* [12]. After the transient increase, IFT88 levels at the base dropped and plateaued at around 70% of the initial intensity values. One possible explanation for this observation is that there is a role for kinesin-2 in temporarily tethering IFT trains to the base of the cilium before their departure. Without functional kinesin-2 motors, the IFT particles detach from the cilium base, resulting in the observed reduction in IFT88 intensity. Alternatively, there may be a feedback mechanism that regulates the amount of IFT protein at the base in response to a cue reporting the status of the cilium [51–53].

### Implications for mechanisms of cilium disassembly

After the block in IFT in response to kinesin-2 inhibition, there was a rapid decrease in the number of ciliated cells, with a nearly complete loss of cilia by six hours of kinesin-2 inhibition. Surprisingly, the average cilium length of the remaining cilia decreased only slowly and moderately over the time course of kinesin-2 inhibition. Interestingly, similar changes in the percentage of ciliated cells and the length of the remaining cilia were observed in *Chlamydomonas* upon a shifting cells of the fla10-1^ts^ strain to the non-permissive temperature [49, 54]. One possibility is that the cilia that persist contain kinesin-2 motors that are not completely inhibited. This seems unlikely, however, because IFT was completely abolished after addition of the B/B inhibitor in every cilium we observed by live-cell imaging.

A second possibility is that there is a two-step mechanism that leads to ciliary disassembly in response to a block in anterograde transport: an initial event that occurs slowly and stochastically and primes the cilium for a subsequent fast disassembly event. The initial event could be the shedding of the tip of the cilium, which has been reported to be a requirement for ciliary disassembly in response to mitogenic signals [55]. The biological purpose of tip shedding could be the efficient and permanent removal of IFT-B from the periciliary environment [55]. In *Chlamydomonas*, the IFT-B returned to the cilium base is available for subsequent rounds of IFT train assembly and injection [12] and thus a loss of IFT-B through tip shedding may prime the cilium for disassembly.

The rapid loss of cilia in response to kinesin-2 inhibition contrasts with the effects of inhibiting dynein function. Treatment of ciliated cells with the small-molecule inhibitor Ciliobrevin D [38, 56] causes IFT to stop within ∼5 minutes [34], however, we observed the first disassembly of cilia 12 hours after Ciliobrevin D treatment and even after 24 hours of treatment, more than 50% of the cells still had a primary cilium, consistent with other reports [38, 57]. *Chlamydomonas* mutants with a temperature-sensitive defect in the dynein-2 motor also showed a dramatic reduction of retrograde IFT yet remained nearly full-length for many hours at the non-permissive temperature [58, 59].

The dramatic requirement for kinesin-2 function in ciliary maintenance is likely because kinesin-2 is required to drive enrichment of cilium-specific proteins against a concentration gradient whereas dynein-2-driven retrograde transport is not the only mechanism by which ciliary proteins can leave this organelle. Recent work in both Chlamydomonas and mammalian primary cilia has demonstrated that various ciliary proteins and motors return to the ciliary base largely by diffusion [60–62], and may provide a mechanism for how cilium length is regulated [63].

Several mechanisms can be envisioned for how cilia disassemble in response to an absence of anterograde IFT. One possibility is that cilium disassembly is a passive process of microtubule disassembly due to the loss of tubulin delivery via IFT. According to the balance-point model, cilium length is determined by an equilibrium of axoneme assembly and disassembly rates [64], and an acute block of tubulin import should result in a continuous decrease in cilium length before the number of ciliated cells is significantly affected. In contrast, we find that cilium disassembly is dominated by a fast disassembly mode with stochastic onset. Thus, a passive process of cilium disassembly appears unlikely. An alternative possibility is that loss of IFT leads to the activation of an intrinsic cilium disassembly pathway. For example, loss of IFT from the cilium may trigger the localization or activation of a microtubule-depolymerizing kinesin such as the kinesin-8 KIF19 [65] or the kinesin-13 KIF2A or KIF24 motors [66, 67]. A cilium disassembly pathway involving polo-like kinase 1 and/or Aurora A [68–70] could also account for the fast disassembly kinetics that we observe. Our ability to acutely block IFT and trigger cilium disassembly will allow us to test these possibilities and to examine whether disassembly of the cilium promotes re-entry of cells into the cell cycle [55].

### KIF3A/KIF3B/KAP is the sole and essential motor for IFT in vertebrates

The engineered i3A/i3B motor allowed us to separate the functions of all kinesin-2 motors (KIF3A/KIF3B/KAP, KIF3A/KIF3C/KAP, and KIF17) in the generation and maintenance of primary cilia in mammalian cells. We find that the heterotrimeric kinesin-2 motor KIF3A/KIF3B/KAP is the sole motor required for ciliogenesis. Expression of neither the heterotrimeric motor KIF3A/KIF3C/KAP nor the homodimeric motor KIF17 was able to rescue ciliogenesis in the absence of heterotrimeric KIF3A/KIF3B/KAP function. This finding is consistent with the fact that loss of KIF17 or KIF3C expression has minimal phenotypes in mouse models [23, 24, 71, 72]. Recent work in zebrafish suggested that the heterotrimeric KIF3A/KIF3C/KAP motor is sufficient to drive IFT in a subset of ciliated cells [40]. This may reflect tissue- or specifies-specific adaptation of the kinesin-2-IFT machinery although the possibility of incomplete morpholino penetrance and kinesin-2 down-regulation in the zebrafish work cannot be ruled out.

The inability of KIF17 to rescue ciliogenesis was surprising as mammalian KIF17 is an active kinesin motor [73] that localizes to mammalian cilia [74], binds to IFT proteins [46, 75], moves in an IFT-like manner in mammalian cilia [45, 76], and influences the ciliary localization of some signaling proteins [39, 77]. We thus considered the possibility that heterotrimeric KIF3A/KIF3B/KAP plays a critical role during cilium assembly, but that KIF17 is able to drive sufficient IFT events for cilium maintenance. However, expression of KIF17 did not delay nor prevent cilium disassembly upon acute KIF3A/KIF3B/KAP inhibition. This finding calls into question a theoretical model that KIF3A/KIF3B/KAP and KIF17 take turns in transporting IFT trains along the axoneme [41]. Rather, we propose that in the vertebrate primary cilium, KIF17 motors behave as a passive IFT cargo and/or as motors driving ciliary transport of non-IFT cargoes (**Fig. 7**). This model would explain why *Kif17* mutant zebrafish display only subtle defects in olfactory cilium structure and a delay in photoreceptor outer segment development [72, 78].

**Figure 7.**
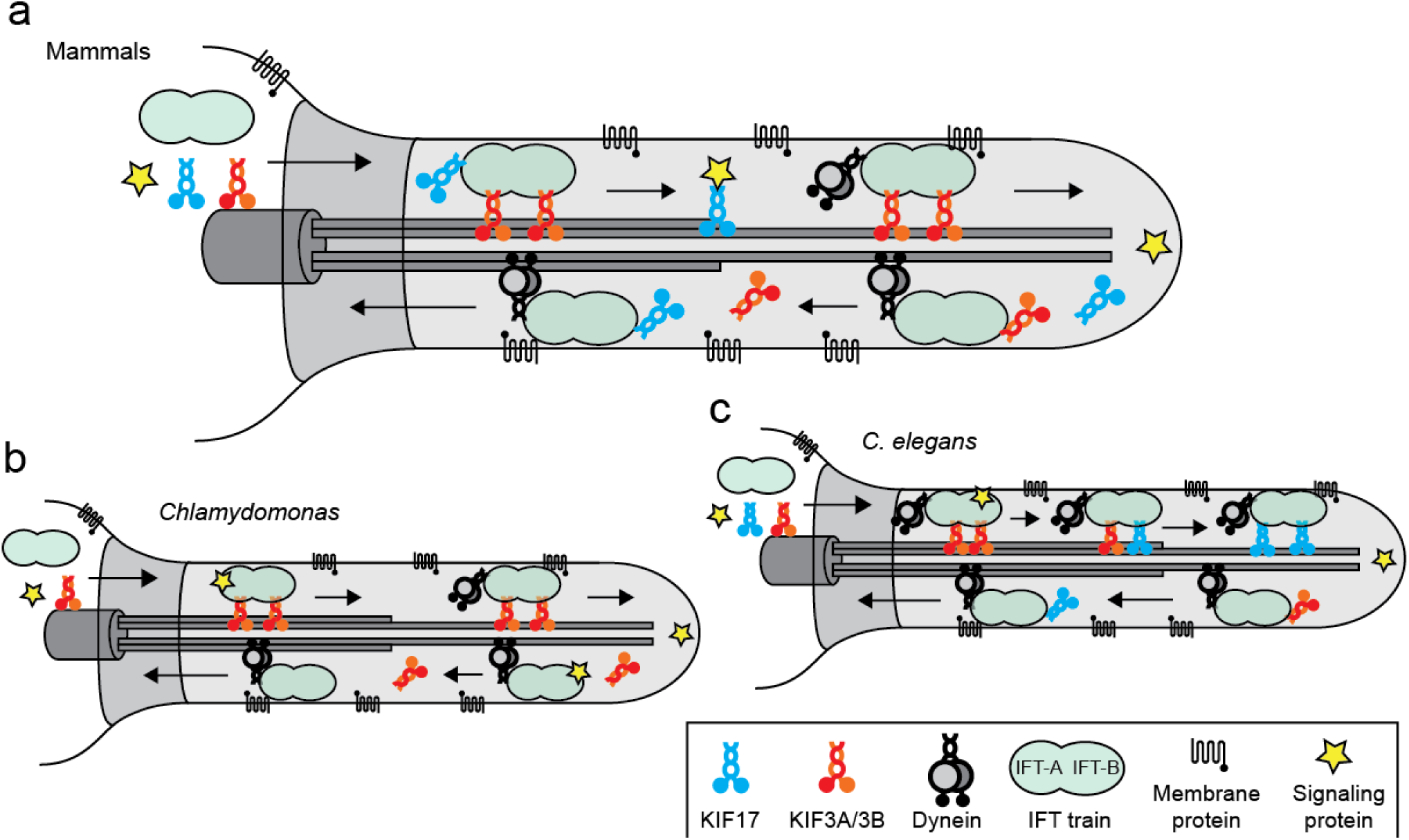
Model of IFT in mammalian cilia. Simplified cartoon representations of IFT in (**a**) mammalian, (**b**) *Chlamydomonas*, and (**c**) *C. elegans* amphid cilia. (**a**) At the base of the cilium, KIF3A/KIF3B/KAP motors bind to preassembled IFT trains for transport along axonemal microtubules. Ciliary proteins and inactive KIF17 and dynein-2 motor proteins are cargoes of the IFT trains. Ciliary membrane proteins track intermittently with IFT. KIF17 cannot drive IFT but could transport other ciliary proteins (e.g. signaling proteins) independently or via its binding to IFT trains. At the tip of the cilium, cargo and motor proteins are unloaded and the IFT trains are transported back to the base of the cilium by cytoplasmic dynein-2. The kinesin-2 motors are returned to the base of the cilium partly by diffusion and in part as passive cargo of IFT trains. (**b**) In *Chlamydomonas*, which lacks a KIF17 homolog, IFT trains are transported in anterograde direction exclusively by heterotrimeric kinesin-II. This motor is returned to the base of the cilium by diffusion. (**c**) In the amphid and phasmid cilia of *C. elegans*, heterotrimeric kinesin-II transports IFT trains through the transition zone. Kinesin-II gradually detaches from IFT trains in the proximal cilium and is successively replaced by the KIF17 homolog, OSM-3, which transports the IFT trains along the distal axoneme. Both kinesin-II and OSM-3 are returned to the base of the cilium by transport as inactive cargo on dynein-driven IFT trains.

### Mechanisms of IFT vary across organisms

We propose a model for how IFT is accomplished in mammalian cilia as compared to models of IFT in other organisms (**Fig. 7**). Current models of IFT are largely based on studies in the ciliated endings of sensory neurons in the amphid and phasmid structures of the nematode *C. elegans*. There, IFT trains are carried along the proximal axoneme by kinesin-II at a slow rate and then handed over to the KIF17 homolog, OSM-3, which solely transports the IFT trains along the distal axoneme at a fast rate ([79], **Fig. 7c**). Genetic and imaging studies have shown that OSM-3 is capable of driving IFT and generating cilia in these sensory structures and that, in contrast, loss of heterotrimeric kinesin-II motor function has only minor effects on cilium structure [16]. These findings provide the basis of current models in the field that KIF3A/KIF3B/KAP is a slow and weak motor and its role is to aid in the navigation of IFT particles through the transition zone [9].

In contrast, our data suggest a model for IFT in mammalian cilia in which the heterotrimeric KIF3A/KIF3B/KAP and homodimeric KIF17 motors play opposite roles from those described in *C. elegans* (**Fig. 7a**). We propose that the heterotrimeric KIF3A/KIF3B/KAP motor is the sole and essential motor for IFT in mammalian cilia and that the homodimeric KIF17 motor plays an accessory role either to transport ciliary cargoes in an IFT-independent manner and/or as an adaptor to link specific ciliary cargoes to the IFT particles. This model for IFT in mammalian cells is similar to mechanisms of IFT transport in *Chlamydomonas* (**Fig. 7b**) where heterotrimeric KIF3A/KIF3B/KAP is the sole and essential motor for IFT transport and a homolog for homodimeric KIF17 has not been found [22]. A separation of function between the heterotrimeric KIF3A/KIF3B/KAP and homodimeric OSM-3 motors has also been noted in the cephalic male (CEM) neurons of *C. elegans* [80].

Species-specific differences are also noted when considering the recycling of the kinesin motors after delivery of IFT particles to the tip of the cilium. In mammalian cilia, kinesin motors have been observed to undergo both retrograde transport and free diffusion back to the cell body [45, 61]. In *Chlamydomonas* cilia, the kinesin motors do not rely on dynein, rather they return to the base of the cilium by diffusion [51, 58, 62, 81]. In contrast, in *C .elegans*, the kinesin-II motors have been observed to undergo retrograde transport and to accumulate at the ciliary tips of dynein-2 mutants [82]. Further work is required to identify the molecular mechanisms that mediate these species-specific differences.

#### Outlook

Future work with the i3A/i3B motor will allow us to directly determine the ciliary cargoes that depend on KIF3A/KIF3B/KAP activity for import into cilia and/or transport along the axoneme. We will also be able to directly test the roles of kinesin-2 in the steps that lead to ciliogenesis, such as the assembly of the subdistal appendages and the positioning of the basal body [83]. Furthermore, the i3A/i3B motor will be a valuable tool to dissect other dynamic cellular processes in which KIF3A/KIF3B/KAP has been implicated, such as neurite outgrowth in neuronal cells [84].

## Methods

### Plasmids and oligonucleotides

For CRISPR/Cas9-mediated genome-editing, the plasmid eSpCas9(1.1) was a gift from Feng Zhang ([85], Addgene plasmid #71814). sgRNAs targeting a site in the third exon of the *Mm Kif3a* gene were generated by annealing the primers 5’-CACCGAAGGCGTTCGAGCAGTACCG-3’ and 5’-AAACCGGTACTGCTCGAACGCCTTC-3’. sgRNAs targeting a site in the second exon of the *Mm Kif3b* gene were generated by annealing the primers 5’-CACCGGTGAAAACCACATCCGGGT-3’ and 5’-AAACACCCGGATGTGGTTTTCACC-3’. The resulting DNA pieces were subcloned via BbsI restriction digestion into eSpCas9(1.1). The plasmid PGK-puro was a generous gift from Dr. Thom Saunders (University of Michigan). Plasmids encoding mCitrine and HsKIF17-mCit [73], mCherry (mChe, Clontech Laboratories), and mNeonGreen (mNG) [86] have been described previously. To generate plasmids encoding full length MmKIF3A-mNG and MmKIF3B-mChe, the open reading frames (ORF) of KIF3A and KIF3B were synthesized (Life Technologies) and cloned in frame via NheI and BamHI into pmNeonGreen-N1 and pmCherry-N1, respectively. The plasmid encoding full length MmKIF3C-mChe was cloned by amplifying the ORF of KIF3C with the primers 5’-GCGTATACTAGTGCCACCATGGCCAGTAAGACCAAGGCCAG-3’ and 5’-CGCATAGGATCCATGTCATGGTCTACCACTGTTGCAGGG-3’ from a plasmid that was a gift from L. Goldstein (University of California at San Diego). The KIF3C ORF was then cloned via the compatible ends of NheI/SpeI and BamHI into pmCherry-N1. Inserts encoding DmrB-KIF3A-mNG, DmrB-Δ11KIF3A-mNG, DmrB-Δ12KIF3A-mNG, DmrB-Δ13KIF3A-mNG, DmrB-KIF3B-mChe, DmrB-Δ5KIF3B-mChe, DmrB-Δ6KIF3B-mChe, and DmrB-Δ7KIF3B-mChe were generated by Splice by Overlap Extension PCR [87, 88] and cloned into pmNeonGreen-N1 and pmCherry-N1, respectively. Primer sequences are found in **Supp Table 1**. To generate the constructs encoding inhibitable motors without a fluorescent tag, the ORFs encoding DmrB-Δ12KIF3A and DmrB-Δ6KIF3B were cloned via NheI and AgeI into a pEGFP-N1 backbone from which the EGFP had been removed. The HH-responsive luciferase encoding plasmid *ptch*Δ136-GL3 [31], pSV-β-galactosidase (Promega, E1081), pCIG-GLI2ΔN [31] were used for luciferase assays. The plasmid IFT88-mNG was cloned by replacing mCit with mNG through restriction digestion with AgeI and BsrGI in the plasmid myc-IFT88-mCit described [89]. All constructs were verified by DNA sequencing.

### Cell culture and genome engineering

Parental NIH-3T3 Flp-In cells (Thermo Fisher Scientific; RRID: CVCL_U422) and derived *Kif3a*^*-/-*^;*Kif3b*^*-/-*^ cells were cultured in D-MEM (Gibco) with 10% Fetal Clone III (HyClone) and GluteMAX (Gibco) at 37°C and 5% CO_2_. Both cell lines were transfected using Lipofectamine 2000 (Life Technologies) according to the manufacturer’s protocol. Double knockout cells (*Kif3a*^*-/-*^;*Kif3b*^*-/-*^) were generated by sequentially engineering the *Kif3a* and *Kif3b* gene loci. NIH-3T3 Flp-In cells were co-transfected with the eSpCas9(1.1) encoding an appropriate sgRNA (see above) and PGK_puro, which confers Puromycin resistance. One day later, 10 μg/ml Puromycin (Sigma, P8833) was added to the culture medium to select for transfected cells. Three days later, single cells were sorted into separate wells of a 96-well plate (Flow Cytometry Core, University of Michigan) and expanded. To identify cell clones with altered KIF3A or KIF3B alleles, the DNA of cells not able to generate cilia in response to serum-starvation was isolated using the DNeasy Blood & Tissue Kit (Qiagen) and the DNA region of interest was amplified by PCR using the primers listed in **Supplementary Table 1.** The PCR product was cloned (TOPO TA Cloning Kit; Invitrogen) and transformed into competent TOP10 bacteria (Thermo Fisher Scientific). Plasmids were isolated (GeneJET Plasmid Miniprep Kit, Thermo Fisher Scientific) and sequenced using the primer 5′-CAGGAAACAGCTATGACC-3′.

### Immunohistochemistry and epifluorescence microscopy

Cells were fixed with 4% formaldehyde in PBS, treated with 50 mM NH_4_Cl in PBS to quench unreacted formaldehyde, permeabilized with 0.2% Triton X-100 in PBS, and blocked in blocking solution (0.2% fish skin gelatin in PBS). Primary and secondary antibodies were applied in blocking solution at room temperature for 1 h each. Cells were incubated 3x for 5 min in blocking solution to remove unbound antibodies. Antibodies used: polyclonal antibodies reacting with ARL13B (1:1000, Protein Tech Group, 17711-1-AP), IFT88 (1:500, Protein Tech Group, 13967-1-AP), and monoclonal antibodies reacting with ARL13B (1:200, NeuroMAB, 73-287), acetylated tubulin (1:10.000, Sigma, T6793), and polyglutamylated tubulin (1:1000, Adipogen Life Sciences, GT335). Nuclei were stained with 10.9 μM 4′,6-diamidino-2-phenylindole (DAPI) and cover glasses were mounted in ProlongGold (Life Technologies). Images were collected on an inverted epifluorescence microscope (Nikon TE2000E) equipped with a 60x, 1.40 numerical aperture (NA) oil-immersion objective and a 1.5x tube lens on a Photometrics CoolSnapHQ camera driven by NIS-Elements (Nikon) software. To measure IFT88 content of cilia, a region of interest (ROI) was drawn around the cilium shaft (identified by ARL13B) and the base of the cilium (identified by glutamylated tubulin) in ImageJ (NIH) and the average fluorescence intensity of IFT88 staining in each of the ROIs was measured and the population average was calculated using Excel.

### Ciliogenesis rescue and disassembly assays

Cells were seeded on cover glasses. Twelve hours later, the culture medium was switched to 1% Fetal Clone III (serum-starvation) and the cells were transfected. Two days later the cells were fixed and stained. Vehicle (0.1% ethanol final) or 50 nM B/B homodimerizer (or as indicated in **Supp Fig. 1**) (Clonetech, 635060) was either added just before the transfection complexes To assess the effect of B/B inhibitor or Ciliobrevin D on fully-formed cilia, cells were seeded on cover glasses and serum-starved (and transfected, B/B inhibitor experiment) twelve hours later. Two days later, vehicle (0.1% ethanol final) or 50 nM B/B homodimerizer or 30 μM Ciliobrevin D (Sigma, 250401) was added for the time spans as indicated in **Fig. 5a,d** and cells were fixed and stained. Image analysis was performed using ImageJ (NIH). Cells expressing mCherry, or mCherry- and mNeonGreen-tagged inhibitable motors were selected and analyzed for the presence of a cilium, as judged by an ARL13B-positive filament that was ≥ 10 pixel (≥1.1 μm). We noted that cells that express low levels of the KIF3B and KIF3A constructs had the highest probability to generate cilia. Thus cells expressing high levels of transfected proteins were excluded from analysis.

### HH-responsive luciferase assay

Cells were seeded onto gelatin-coated 24-well plates (5 x 10^4^ cells/well). Twelve hours later, cells were co-transfected with WT or i3A/i3B constructs, a luciferase reporter (*ptch*Δ136-GL3), and β-galactosidase control (pSV-β-galactosidase), or were co-transfected with i3A/i3B constructs and pCIG-GLI2ΔN, a luciferase reporter (*ptch*Δ136-GL3), and β-galactosidase control (pSV-β-galactosidase). Two days after transfection, cells were briefly washed with PBS and transferred to culture medium containing 1% Fetal Clone III, 1% penicillin/streptomycin and 50 nM B/B inhibitor and 500 nM SAG (Enzo Life Sciences, ALX-270-426-M001), as indicated. Luciferase and β-galactosidase activities were measured after two days of serum starvation using a Spectra-max M5e plate reader (Molecular Devices) and the luciferase assay kit (Promega, E1501) and β-galactosidase assay kit (EMD Millipore, 70979). To calculate the level of HH pathway activity luciferase values were divided by the β-galactosidase values, and data were reported as fold induction relative to either vector transfected or untreated values. All treatments were performed in triplicate.

### Live-cell imaging

Cells were seeded into glass bottom dishes (MatTek Corporation, P35G-1.5-14-C) and 12 hours later switched to culture medium containing 1% serum and transfected. Two days later, medium was changed to Leibovitz’s L-15 Medium (Gibco, 21083-027) and imaged on a Nikon Ti-E/B microscope equipped with a 100x, 1.49 NA oil-immersion TIRF objective, 1.5x tube lens, three 20mW diode lasers (488, 561, 640 nm) controlled via acousto-optical tunable filter (AOTF) (Agilent) and EMCCD detector (iXon X3 DU897, Andor). Time-lapse images were acquired with 488 nm excitation, 200 ms exposure at 5 or 2.5 frames per second. Kymographs were generated using the Multiple Kymograph plugin (J. Rietdorf and A. Seitz) for ImageJ (NIH).

### Statistical analysis

Statistical tests were performed in the software GraphPad Prism (Version 7, GraphPad Software). The statistical significance between different treatments was first tested via one-way ANOVAs: Fig. 3 (p < 0.0001), Fig. 4c (Kruskal-Wallis nonparametric, p < 0.001), Fig. 5b (p < 0.001), Fig. 5e (p = 0.0053). For Fig. 6 a two-way ANOVA found a significant effect of treatment time (Fig. 6b, p<0.0001; Fig. 6c, p = 0.0006) but no effect of the expressed protein (mCit vs. KIF17) and no interaction between the treatment time and the expressed protein. Post hoc statistical tests were then applied as described in each figure legend.

## AUTHOR CONTRIBUTIONS

K.J.V., M.F.E., B.W., B.L.A designed the experiments. M.F.E., B.W., S.E.K. and A.S. performed experiments. M.F.E., B.W., S.E.K., A.S., and A.B. analyzed the data. K.J.V. and M.F.E. wrote the manuscript, with input from all authors.

## ACKNOWLEDGMENTS

This work was supported by awards from the National Institute of General Medical Sciences (R01GM116204 to K.J.V., R01GM118751 to K.J.V. and B.L.A.), the National Cancer Institute (R01CA198074 to B.L.A.), and the National Institute on Deafness and other Communication Disorders (R01DC014428 to B.L.A.) of the National Institutes of Health. We gratefully acknowledge T. Saunders and J. Adams (University of Michigan) for support with generating knockout cell lines and statistical analysis, respectively.

## SUPPLEMENTARY MATERIAL

**Supplementary Table 1.**
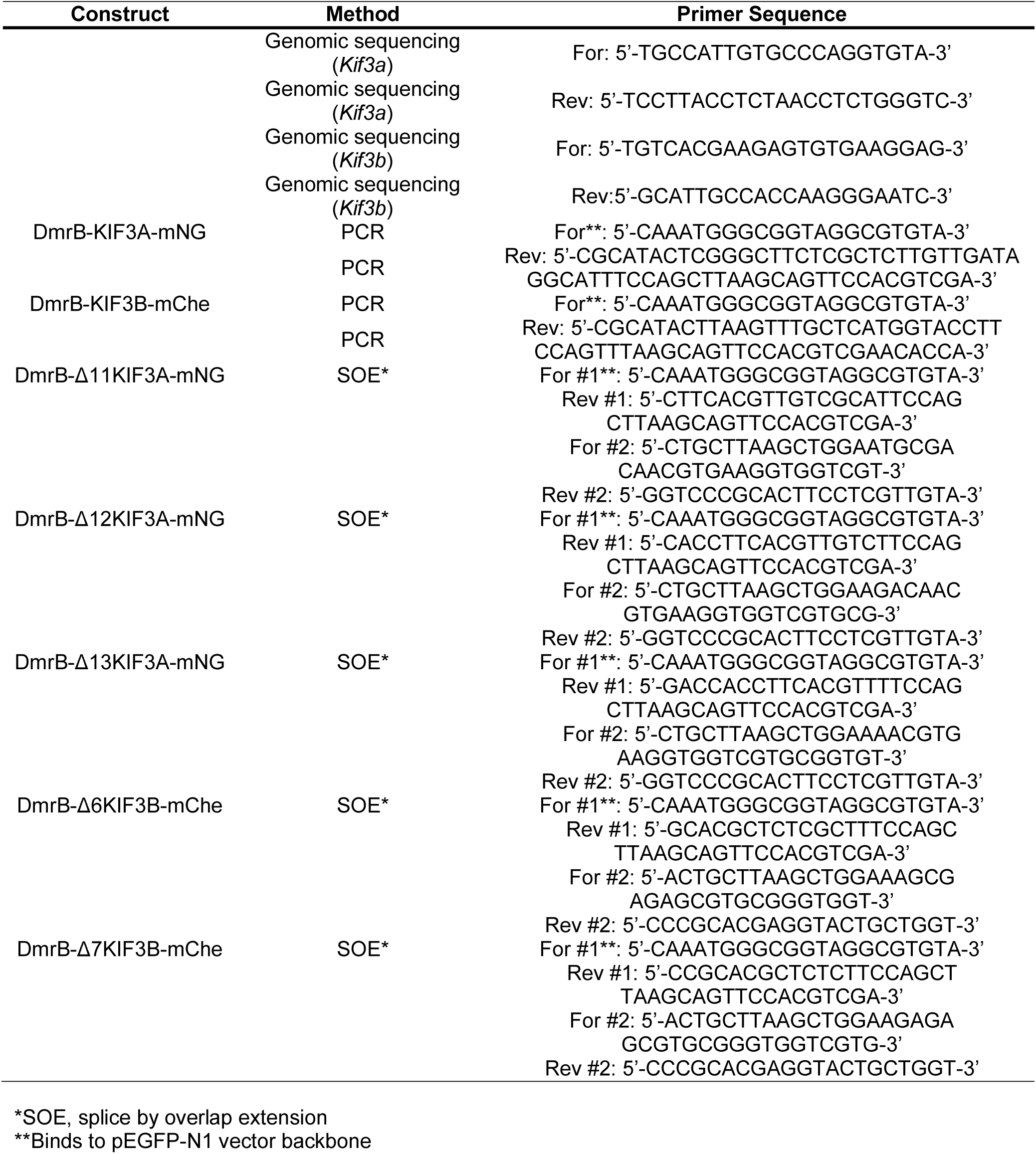
Primer sequences.

**Supp Fig 1.**
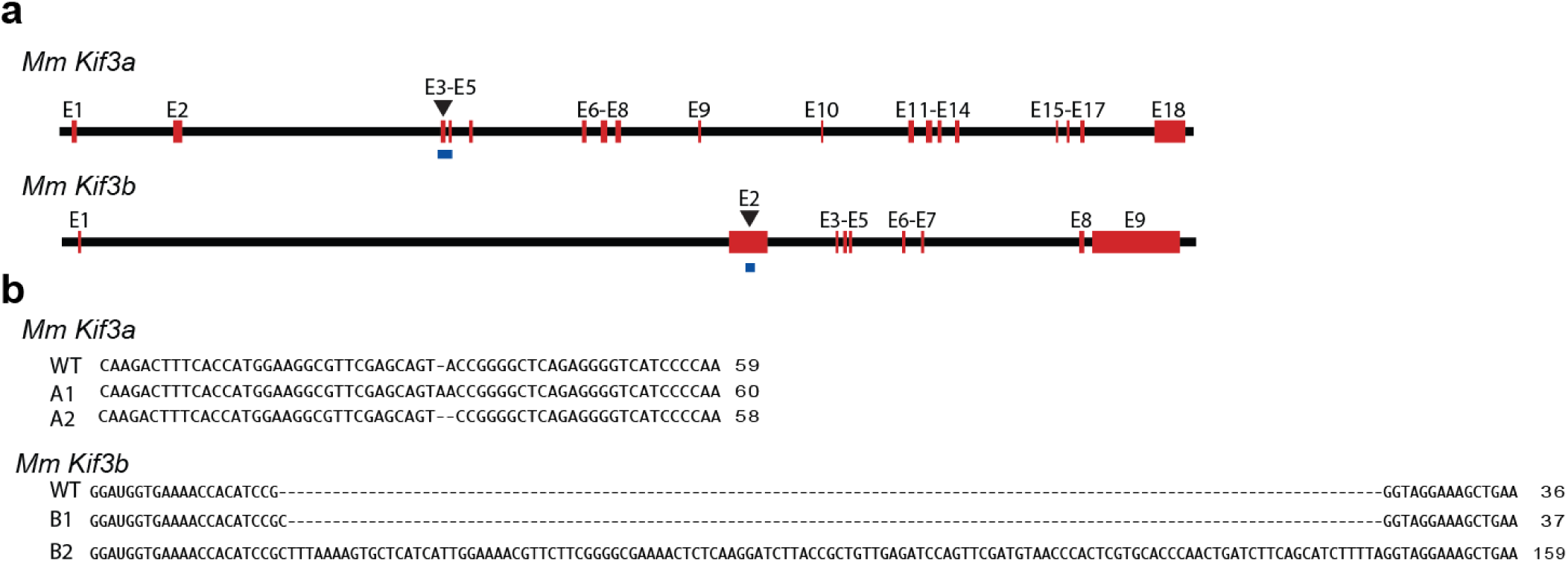
Generation of KIF3A/KIF3B double knockout NIH-3T3 cells using CRISPR/Cas9. (**a**) Map of the murine genes for *Kif3a* (chromosome 11) and *Kif3b* (chromosome 2). Exons (E) are labelled in red, the binding site of the single guide RNA used to generate knockout cells via CRISPR/Cas9 genome editing is indicated with a black triangle, and the PCR product generated for assessment of CRISPR/Cas9 cutting is shown in blue. (**b**) After successive editing of KIF3A and then KIF3B, a cell clone was subjected to DNA sequencing. Ten Topo-cloned PCR fragments revealed only two allele variants for *Kif3a* (A1 and A2) and two allele variants for *Kif3b* (B1 and B2), confirming that the genetically-engineered cell line is clonal. In comparison with the KIF3A WT sequence, allele A1 has a single base-pair (bp) insertion and allele A2 has a 1 bp deletion, both of which lead to a frameshift that results in a premature stop codon in E4. In comparison with the KIF3B WT sequence, allele B1 has a 1 bp insertion and allele B2 has a 123 bp insertion, both of which lead to a premature stop codon in E2. All potentially translated proteins are shorter than 278 amino acids and thus would not encode the kinesin motor domain of ∼340 amino acids.

**Supp Fig 2.**
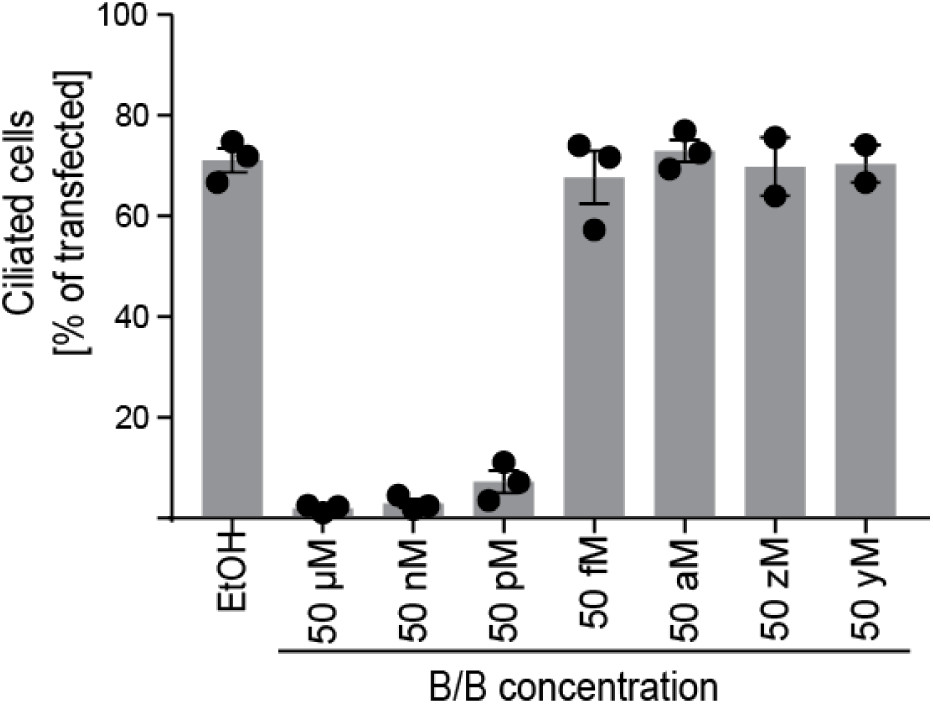
B/B inhibitor dose-response curve. *Kif3a*^*-/-*^;*Kif3b*^*-/-*^ cells were transfected with inhibitable kinesin-2 (i3A^Δ12^ and i3B^Δ6^) motors and serum-starved for two days in the presence of ethanol vehicle (EtOH) or different concentrations of B/B inhibitor, as indicated. The percentage of cells expressing both i3A^Δ12^ and i3B^Δ6^ subunits that contain a primary cilium was quantified and the means of 2-3 independent experiments are shown. Error bars, SEM.

**Supp Fig 3.**
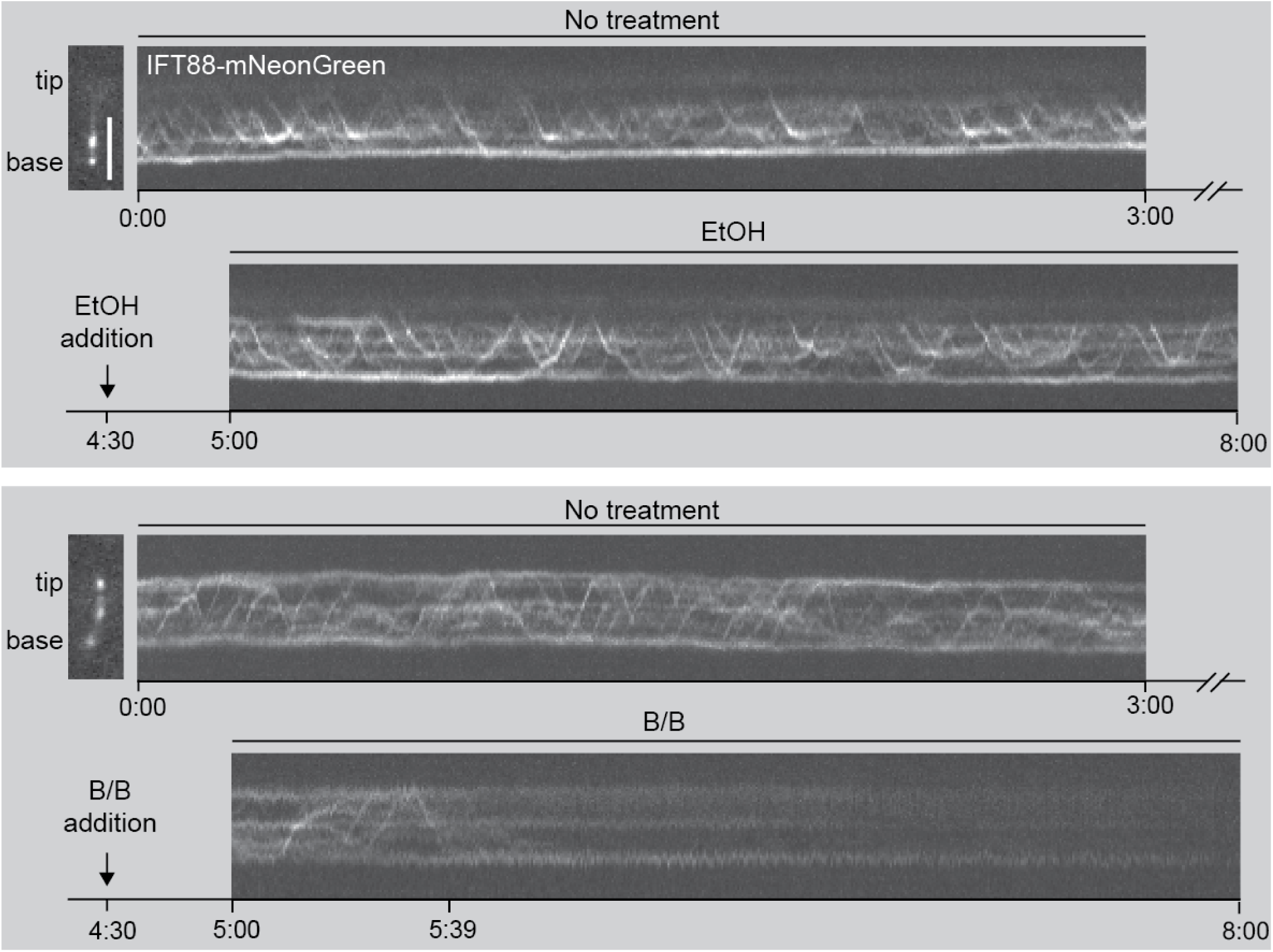
Kinesin-2 inhibition abruptly stalls IFT. Cells were processed and image as described for **Fig. 4**, but imaging frequency was adjusted to 2.5 Hz in order to extend the total imaging interval. Kymographs from two representative cilia are shown. The upper panel shows a cilium of a control treated (EtOH) cell and the lower panel shows a cilium of a cell treated with 50 nM inhibitor (B/B). IFT motion ceases 1:09 minutes after B/B inhibitor addition.

